# *In-Silico* Investigation of Luminol, Its Analogues and Improved Mechanism of Chemiluminescence for Blood Identification Beyond Forensics

**DOI:** 10.1101/2020.01.08.899617

**Authors:** Toluwase Hezekiah Fatoki

## Abstract

This study aimed to discover chemiluminescent analogues of luminol, understand their molecular binding to hemoglobin of bloodstains in the household crime, and the mechanism of chemiluminescence. Similarity and clustering analyses of luminol analogues were conducted, and molecular docking was carried out on hemoglobin from *Homo sapiens* and other four domestic organism namely *Gallus gallus, Drosophila melanogaster, Rattus norvegicus,* and *Canis familiaris*. The results show that the order of overall binding score is *D. melanogaster > H. sapiens > C. familiaris > R. norvegicus > G. gallus*. Seven compounds namely ZINC16958228, ZINC17023010, ZINC19915427, ZINC34928954, ZINC19915369, ZINC19915444, and ZINC82294978, were found to be consistently stable in binding to diverse hemoglobin and possibly have chemiluminescence than luminol. The amino acid residues involved in the interaction of human hemoglobin with the 30 test compounds, show that His45, Lys61, Asn68, Val73, Met76, Pro77, Ala79, Ala82, Leu83, Pro95, Phe98, Lys99, Ser102, Ser133, Ala134, and Thr134 are significant in the mechanism of action of presumptive test compounds. The improved mechanism of chemiluminescent identification of blood hypothesized that nitrite interact with the Fe(II) heme, with the cleavage of a hydroxide ion and the formation of the nitrosonium cation in peroxidase reaction. It was proposed that degradation of rhombic heme complex to fluorescent products is possibly inhibited by nitric oxide from the test compound luminol. This study provides novel insight on the luminol and its actual mechanism for broader possible applications of luminol with careful development of new methodologies.

## INTRODUCTION

Forensic biology is an aspect of forensic science that comprises of serology and deoxyribonucleic acid (DNA) analyses. Forensic science is the application of science to the criminal and civil laws that are enforced by police agencies in a criminal justice system (Saferstein, 2011; Fatoki, 2016). In the forensic investigation laboratory, serology analysis deals with the screening of evidence for the presence of body fluids (such as bloodstains, saliva and semen) while DNA profiling is used to pinpoint body fluids to a specific organism or living being. However, the strengths and present limitations of DNA profiling have been presented (Fatoki, 2016).

Blood is an important body fluid that is required for living systemic activities and consists of many components (cells, hemoglobin, proteins antibody, clotting factor etc), useful for metabolic profiling in relevance to forensic toxicology and forensic pathology. The identification of blood is central to many homicide, aggravated assault, sexual assault, and burglary investigations. The presence of blood on evidentiary items can be critical in establishing guilt or innocence during criminal proceedings while blood spatter interpretation provides the manner in which blood was deposited (Gefrides and Welch, 2011). Blood as a crime evidence can be identified through reagents that act on hemoglobin and its derivatives (proteins and amino acids). Due to its immutable and nontransferable characteristics, these components are capable of identifying the origin of evidence (da Silva, 2012).

Luminescence refers to the emission light which occurs when a molecule in an excited state relaxes to its ground state and various types of luminescence differ from the source of energy to obtain the excited state (Dodeigne *et al*., 2000). In chemiluminescence, the energy is produced by a chemical reaction. Luminol (5-amino-2,3-dihydro-1,4-phthalazinedione), is a chemical that has found application in forensic, environmental, biomedical, and clinical sciences (Khan *et al*., 2014). Luminol is used most commonly at crime scenes to reveal bloodstains that are not readily, apparent to the naked eye. The chemiluminescence of luminol is catalyzed by the haemoglobin in blood and it has been used as a very sensitive forensic test for blood (Quickenden *et al*., 2004; Arruda-Vasconcelos *et al*., 2017). Luminol can react vigorously with selected classes of reagents which include strong reducing agents, strong oxidizing agents, strong acids, and strong bases. Report has shown that in aprotic media (dimethylsulphoxide or dimethylformamide), only oxygen and a strong base are required for luminol chemiluminescence while in protic solvents (water, water solvent mixtures or lower alcohols), various oxygen derivatives (molecular oxygen, peroxides, superoxide anion) can oxidize luminol derivatives but required catalyst either enzymes or by mineral catalysts (Dodeigne *et al*., 2000).

Most laboratories also use phenolphthalein, tolidine, tetramethylbenzidine, or leucomalachite green for preliminary blood identification testing (Gaensslen, 2000). The derivatives of isoluminol has been found to reactive to some extent include aminoethylethylisoluminol (AEEI), aminobutylethylisoluminol (ABEI) and aminobutylethylnaphthalhydrazide (ABENH) (Dodeigne *et al*., 2000). Previous research has shown that luminol did not adversely affect the polymerase chain reaction (PCR) during DNA analysis of the bloodstain treated with luminol and that luminol did not interfere with the phenolphthalin and tetramethylbenzidine presumptive tests for blood (Gross *et al*., 1999; Lee *et al*., 1989; Hochmeister *et al*., 1991). However, the major drawbacks to blood detection using luminol are the false-positive results, the lack of specificity, and dark environment conditions (Stoica *et al*., 2016). Thus, there is a need to identify more stable chemiluminescence compounds.

Although, bioinformatic workflows and methods used in other fields of DNA science has been applied to forensic DNA profiling from high-throughput sequencing (Liu and Harbison, 2018) and forensic reconstruction of high-throughput biological assays (Baggerly and Coombes, 2009), the *in silico* serological analysis for forensic science application has not been investigated. The aim of this study is to discover chemiluminescent analogues of luminol, predict their molecular binding to hemoglobin of bloodstains in the household crime, and the mechanism of chemiluminescence.

## MATERIALS AND METHODS

### Similarity search and clustering analysis

The structure of luminol was obtained from the PubChem Compound Database in canonical SMILES (Simplified Molecular Input Line Entry Specification) format and was used to query the ZincDrugLike database using Swisssimilarity server (Zoete *et al*., 2016), to obtained structurally similar compounds (ligands), and they were manually screen for the presence of two ketone functional groups at a similarity score cutoff of 0.800. The structure of other six compounds for blood test were also obtained, namely; phenolphthalein, O-tolidine, tetramethylbenzidine (TMB), leucomalachite green (LMG), aminoethylethylisoluminol (AEEI) and aminobutylethylisoluminol (ABEI). Clustering analysis was performed using ChemMine tools (http://chemmine.ucr.edu/) (Tyler *et al.,* 2011)

### Ligand and target protein preparation

The structure of 25 chemical compounds obtained from similarity search were extracted from the result of SwissSimilarity analysis in SMILES format and converted to MOL2 format using ChemSketch (ACDLabware v2015). The MOL2 files were then converted to protein data bank (pdb) format using PyMol v2.0.7. The crystal structure of *Homo sapiens* (Human) hemoglobin (Hb) and Hb of four other domestic organisms (*Gallus gallus, Drosophila melanogaster, Rattus norvegicus,* and *Canis familiaris*), were obtained from the RSCB Protein Data Bank (www.rscb.org).

### Molecular Docking Studies

The molecular docking studies were carried out according to the method of Fatoki *et al*. (2018). Briefly, all water molecules, hetero atoms, and multichain were removed from the crystal structure of the prepared targets using PyMol v2.0.7. The Gasteiger partial charges were added to the ligand atoms prior to docking. The docking parameter (Table 1) were set up using AutoDock Tools (ADT) v1.5.6 (Morris *et al.,* 2009) and saves the output file of each prepared ligand and each prepared target as pdbqt format. Molecular docking program AutoDock Vina v1.1.2 (Trott and Olson, 2010) was employed to perform the docking experiment from the command line. After docking, the ligands were analyzed and visualized using ADT and PyMol v2.0.7.

**Table 1:** Docking parameter for Hemoglobin of Human and other four domestic organisms.

| SN | Hemoglobin | Center grid box (points) | Size (points) | Spacing (Å) |
| --- | --- | --- | --- | --- |
| 1 | *Homo sapiens*<br>(PDB ID: 1A3N, Chain A) | 5.852 x 12.788 x 21.724 | 120 x 100 x 106 | 0.375 |
| 2 | *Gallus gallus*<br>(PDB ID: 1HBR, Chain A) | 10.250 x 4.361 x 0.514 | 120 x 126 x 104 | 0.375 |
| 3 | *Drosophila melanogaster*<br>(PDB ID: 2BK9, Chain A) | 2.410 x -1.515 x 7.438 | 126 x 106 x 110 | 0.375 |
| 4 | *Rattus norvegicus*<br>(PDB ID: 3DHT, Chain A) | 30.555 x -22.064 x 42.068 | 120 x 108 x 110 | 0.375 |
| 5 | *Canis familiaris*<br>(PDB ID: 3GOU, Chain A) | -16.007 x -7.061 x -3.860 | 116 x 108 x 110 | 0.375 |

### Novel insight on mechanism of action of luminol

The results the previous steps were integrated with available information in available literature through text mining (da Silva *et al*., 2012; Albertin *et al*., 1998; Furniss *et al.*, 1989). The mechanism was fine-tuned with existing organic chemistry reactions, and the new mechanism of chemiluminescence of luminol was proposed.

## RESULTS AND DISCUSSION

Crime scene in a vegetative, down-town and rural evironment pose problems with artifacts which are not readily indistinguishable from human bloodstains and most other body fluids. Domestic enviroment often features birds, rodents, flies and dog. This neccessitate the need for test compounds that can readily specific to human during crime scene sample analyses.

Clustering of compounds by structural or property similarity can be a powerful approach to correlating compound features with biological activity and it is widely used method for diversity analyses to identify structural redundancies and other biases in compound libraries (Sanni *et al.,* 2017). The result of multi-dimensional scaling (MDS) clustering method provided by ChemMine shows that luminol has chemical fingerprint that differes from other commonly used compounds for presumptiv test for blood. All of the identified analogs of luminol except AEEI (compound 6) were found in different cluster (Figure 1), while ABEI (compound 7) and ZINC12427759 (compound 17).

**Figure 1:**
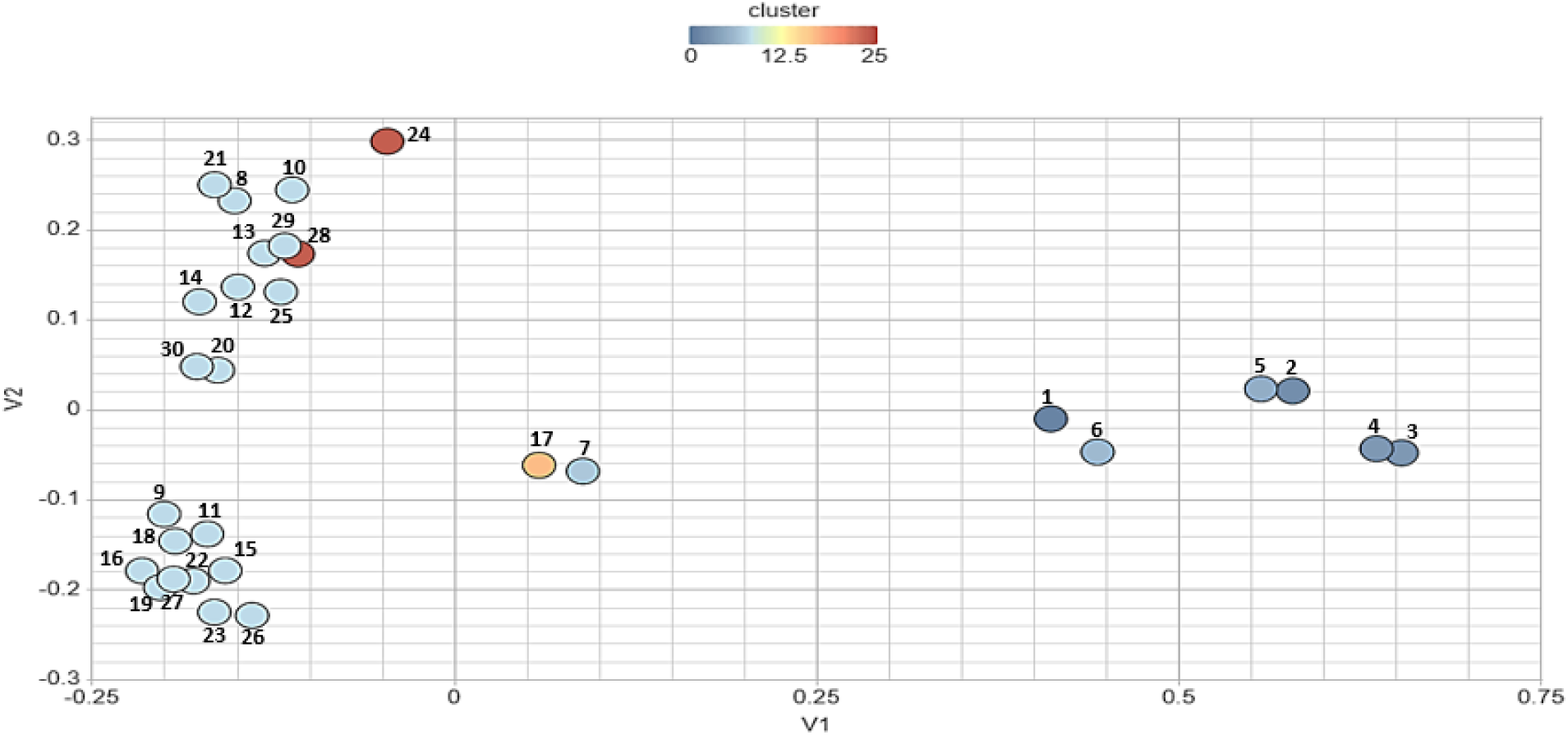
Multi-dimensional scaling clustering (Similarity Cutoff: 0.5, Dimensions: 2D). details of compounds 1-30 are listed in Table 2.

Hemoglobin from *Homo sapiens* and other four domestic organism namely *Gallus gallus, Drosophila melanogaster, Rattus norvegicus*, and *Canis familiaris*, were investigated for their binding to luminol, its analogs and other six compounds used in presumptive test for blood (Table 1 and 2). From the docking result, seven compounds namely ZINC16958228, ZINC17023010, ZINC19915427, ZINC34928954, ZINC19915369, ZINC19915444, and ZINC82294978, were found to be consistently stable in binding to diverse hemoglobin and possibly have chemiluminescence than luminol (Figure 2). The order of overall binding score is *D. melanogaster > H. sapiens > C. familiaris > R. norvegicus > G. gallus*. This shows that *D. melanogaster* can be used as a model organism for forensic research studies without any barrier of ethical permission that often encountered when using human samples.

**Table 2:**
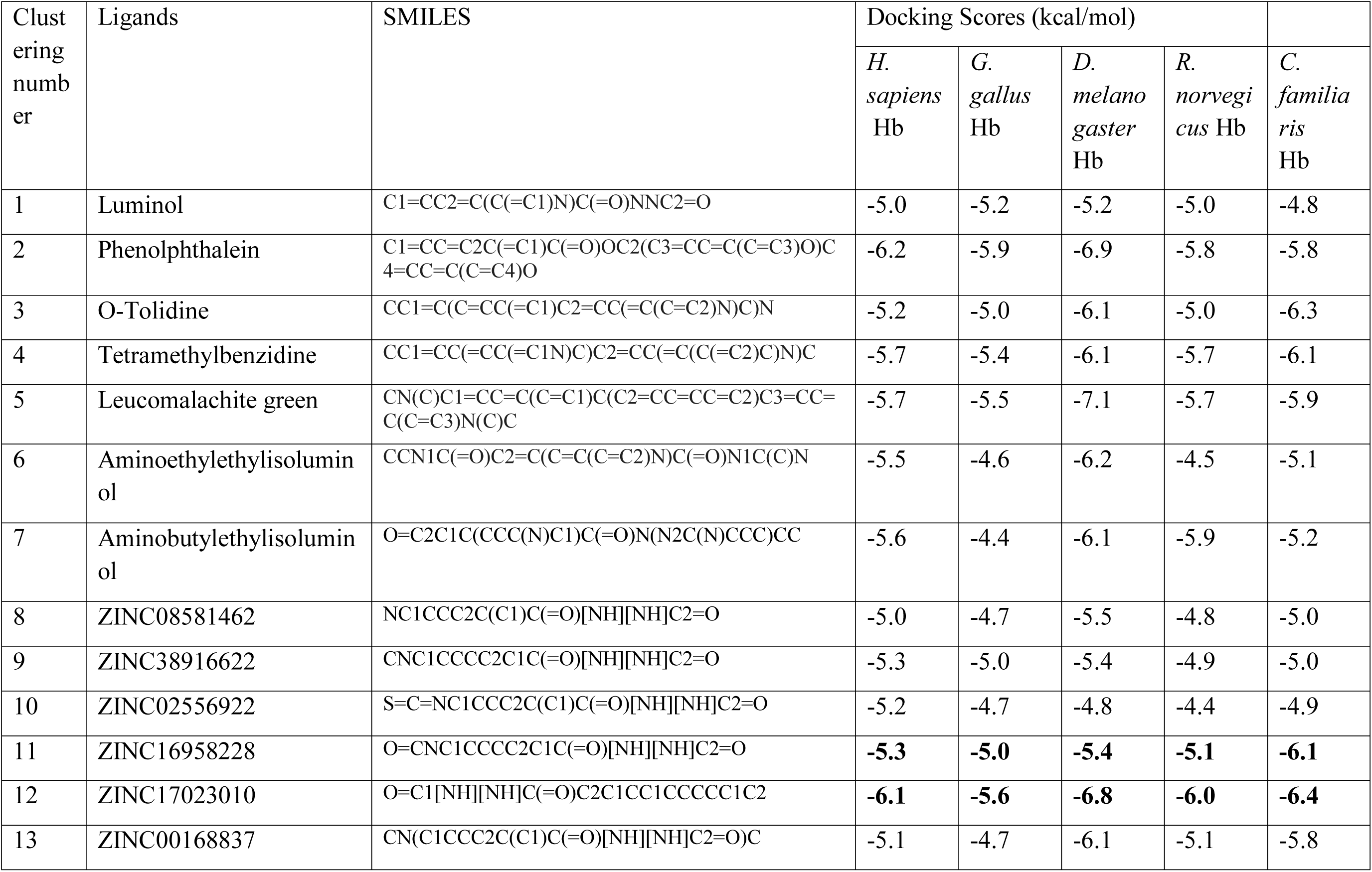
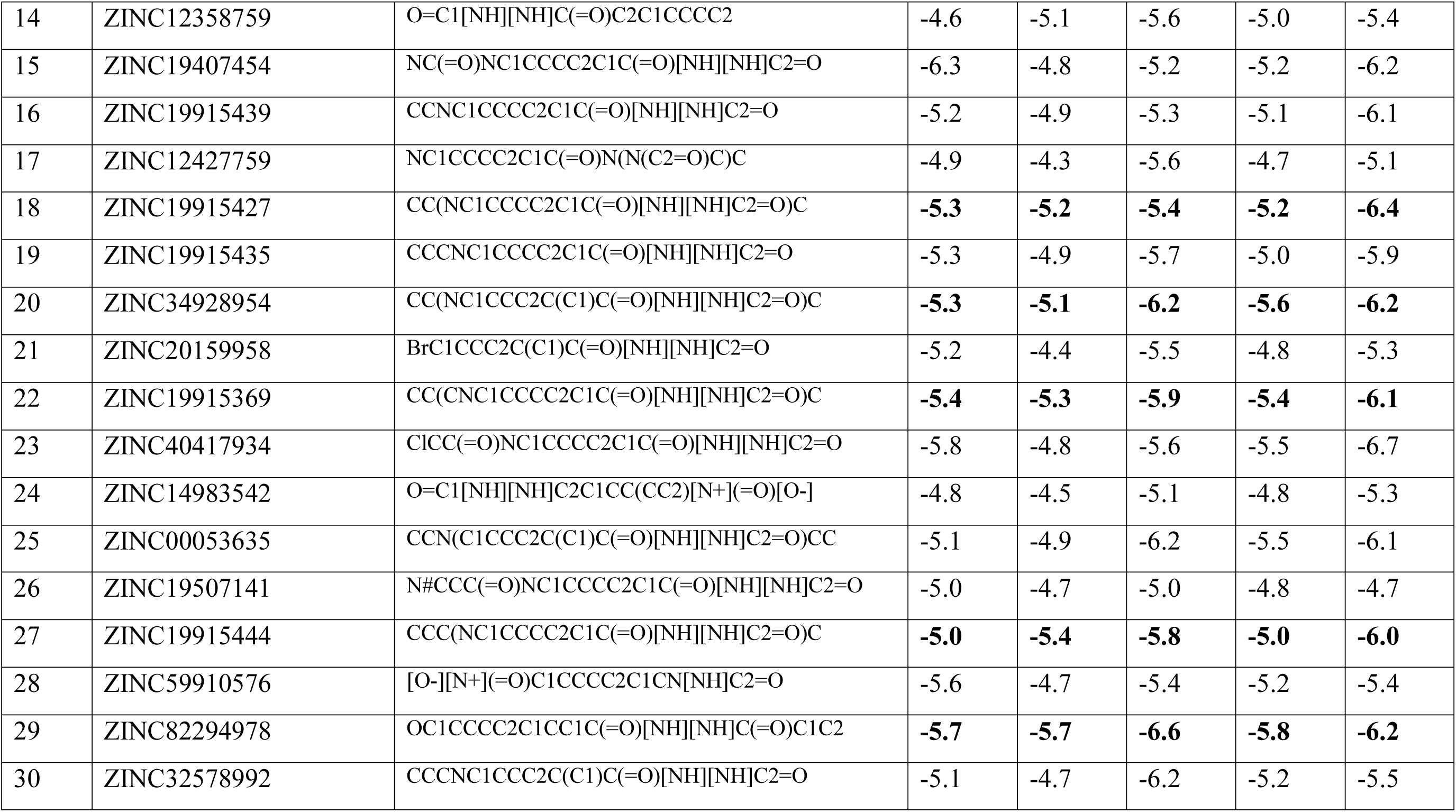
Molecular docking score for luminol, analogs of luminol (zinc compounds) and other compounds used in presumptive test for blood.

**Figure 2:**
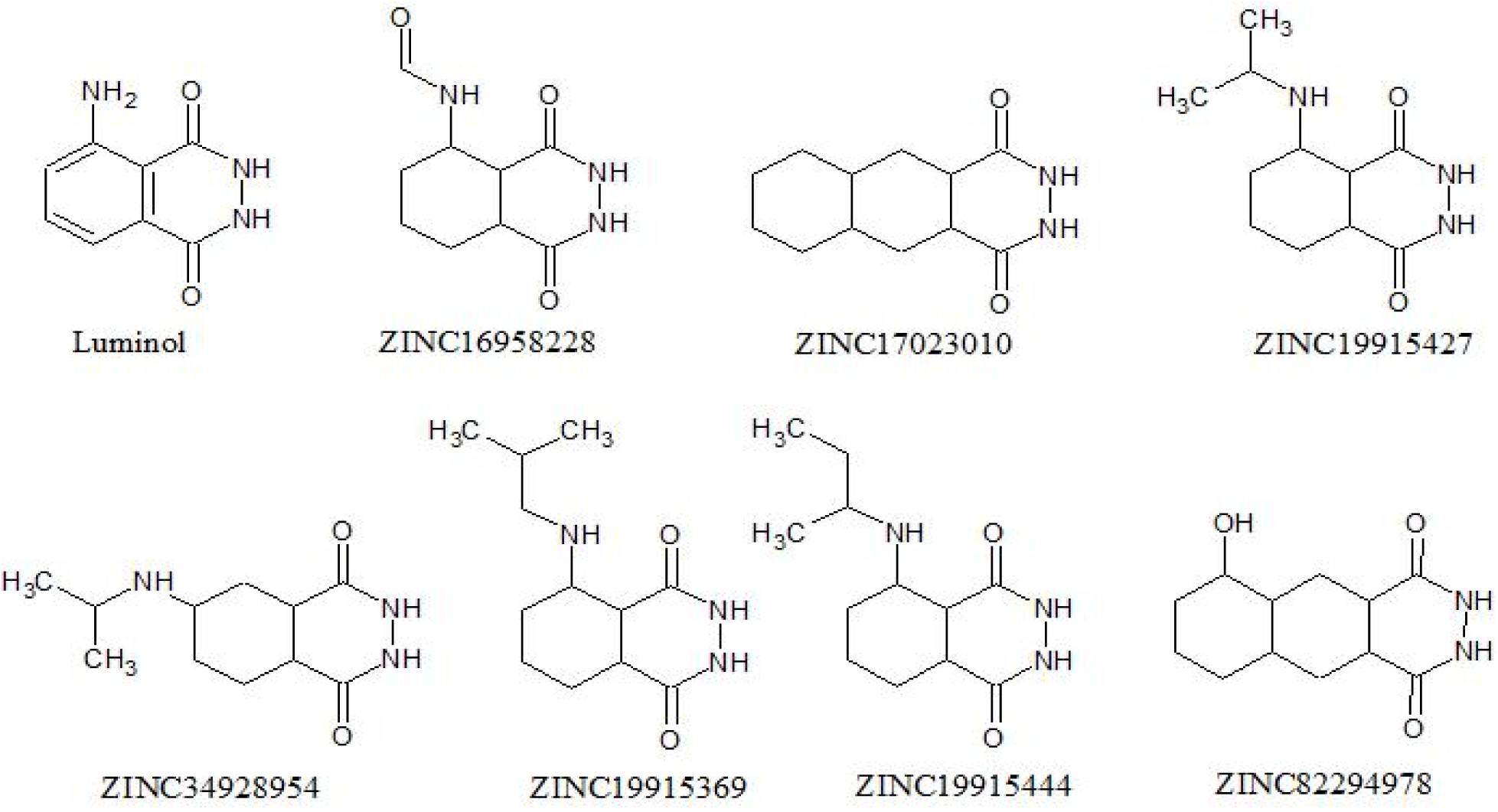
Structure of luminol and its seven predicted stable chemiluminescence analogues.

The Scientific Working Group for Bloodstain Pattern Analysis (SWGSTAIN) has defined insect stains as those bloodstains produced as a result of insect activity (SWGSTAIN, 2008). *D. melanogaster* has been used as a model system for toxicological studies and diseases mechanism such as neurological disorders, cancer, developmental disorders, metabolic and storage disorders and cardiovascular disease (Bier, 2005; Abolaji *et al*., 2014; Saraiva *et al*., 2018). In forensic, *D. melanogaster* and few other insects provide insights to the degradation state of the body fluid in the crime scene (Leitch *et al*., 2018; Jiang and Edgar, 2011, Kulstein *et al*., 2010) and also serve as source of artifacts (Rivers and Geiman, 2017).

The ZINC17023010 (compound 12) which is known as benzo[g]phthalazine-1,4 (2H,3H)-dione, was found to produce chemiluminescence by reaction with hydrogen peroxide in the presence of potassium hexacyanoferrate (III) in an alkaline medium, with low relative chemiluminescence intensities (RCI) than luminol and high relative fluorescence intensities (RFI) than luminol after the chemiluminescence reaction (Yoshida *et al*., 1999). The fluorescence excitation (Ex) and emission (Em) maxima of ZINC17023010 are 360 nm and 415 nm respectively, and thus to generated chemiluminescence at 415 nm (Yoshida *et al*., 1999). Insightfully, from the clustering and docking results of compound 12, it could be hypothesized that compounds 20 and 29, which are among selected compounds broad-spectrum binding affinity for hemoglobin (Figure 2), would have better chemiluminescent properties. This confirmed the probability that the analogs of luminol identified in this study might be utilize as chemiluminescence labelling reagents. As it is summarized in Figure 3, the amino acid residues involved in the interaction of human hemoglobin with the 30 test compounds, show that His45, Lys61, Asn68, Val73, Met76, Pro77, Ala79, Ala82, Leu83, Pro95, Phe98, Lys99, Ser102, Ser133, Ala134, and Thr134 are significant in the mechanism of oxidation of presumptive test compounds.

**Figure 3:**
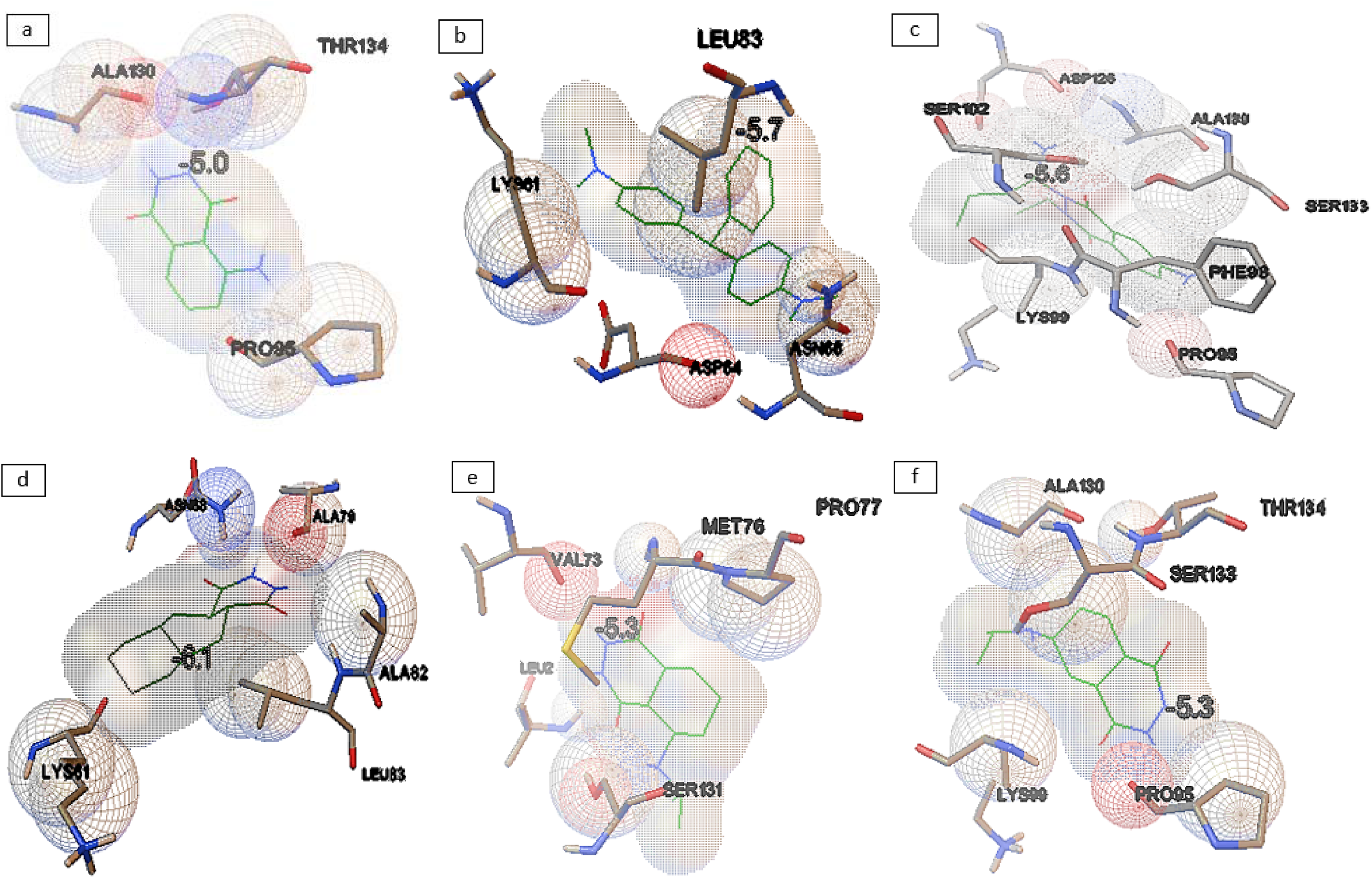
Interaction of human hemoglobin (PDB ID: 1A3N) with (a) Luminol, (b) Leucomalachite green, (c) Aminobutylethylisoluminol, (d) ZINC17023010, (e) ZINC19915427, and (f) ZINC34928954.

The mechanism of hemoglobin (Hb) and its blood derivatives to enhance the oxidation of luminol in the presence of an alkaline solution, is based on the peroxidase-like activity of hemoglobin (An *et al*., 2012). However, studied have attempted to improve the reaction of luminol from Weber protocol (Weber, 1996) through the use of natural cyclodextrin and urea (Stoica *et al*., 2016; Maeztu *et al*., 2010). Stoica *et al*. (2016) investigated the influence of 8 M urea pretreatment on the Weber protocol by molecular dynamic simulation and they observed significantly stronger chemiluminescence signal with almost no false-positive reaction.

What is the impact of NaOH and H_2_O_2_on the heme oxidation? What is the contribution of test compound (such as luminol) and heme on the color of emitted light? The answers to these questions actually lead to the modification of the existing mechanism of action of luminol provided in this study as shown in Figure 4. Majority of the study on hemoglobin have been in relation to its function as oxygen and carbon (iv) oxide carrier in cellular respiratory process. However, in forensic science, actual mechanism of heme in hemoglobin of the red blood cell in identification of blood as crime sample has not been resolved. A comprehensive study has shown that the oxidation mechanism for luminol is still not completely known due to uncertainty which underlies reaction intermediates (Menezes, 2010). The gap in knowledge could be traced to an oversight in taking into cognizance that heme in the hemoglobin is not just an iron (II) ion but a biological multi-oxidative state iron complex that can be degraded by oxidation. Heme comprise of iron (II) which binds to the center of the porphyrin IX by covalent and coordinate bonding (da Silva *et al*., 2012). Study has shown that peroxidase-like activity of heme-like compounds called Prussian blue nanoparticles (PBNPs) was higher than that of Fe (III) and that the catalytic capacity of PBNPs did not simply originate from the Fenton reaction (Chen *et al*., 2018). In addition, the PBNPs were also found to possess catalase-like activity, with the catalytic decomposition of H_2_O_2_ into H_2_O and O_2_ (Zhang *et al*., 2016). The result of the study by Chen *et al*. (2018) implicated the possible presence of Fe(IV)=O species in the FeN*x* unit assembled by C≡N ligands.

**Figure 4:**
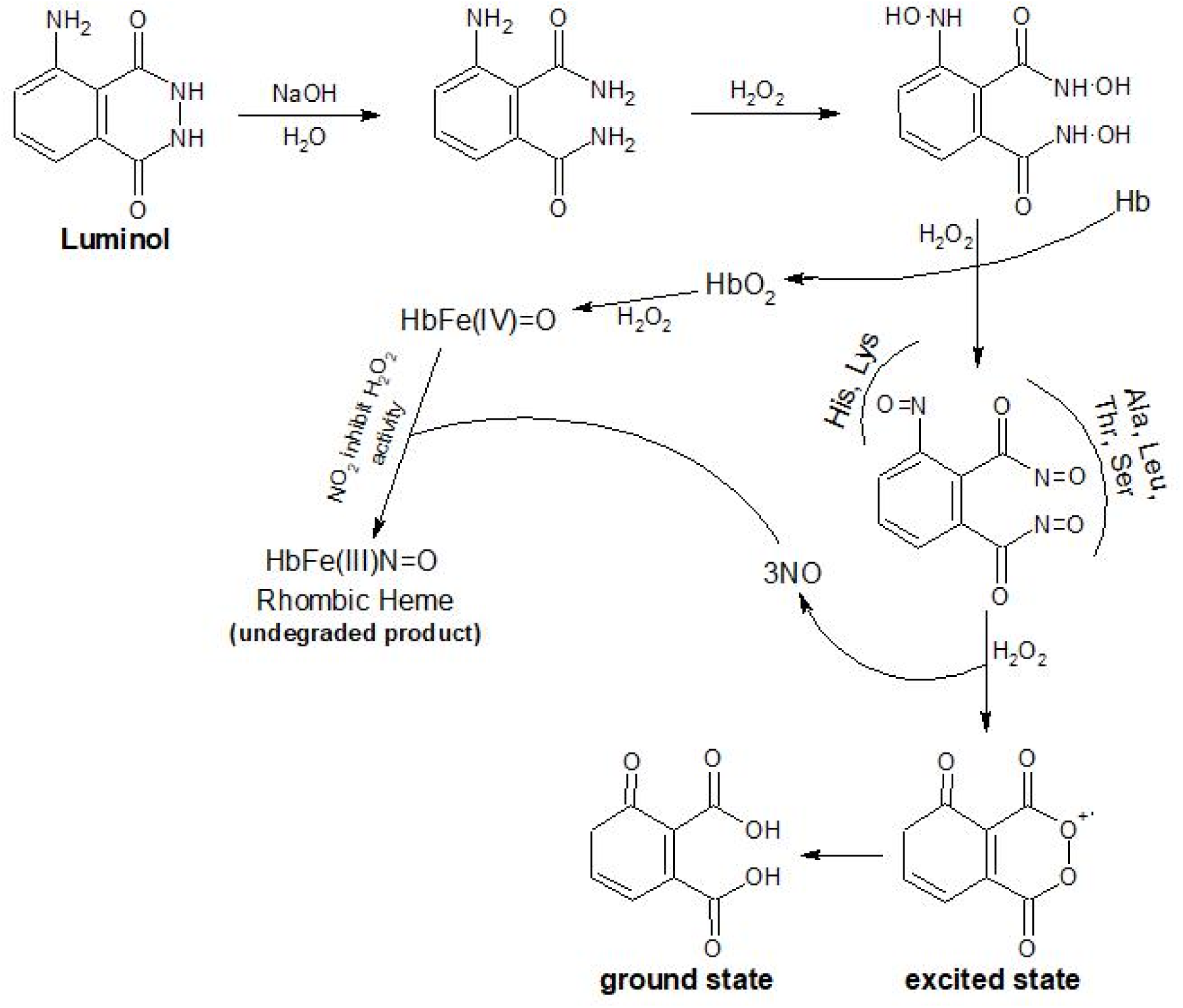
New hypothetical *in-vitro* mechanism of chemiluminescent identification of blood with luminol

Report of the mechanism of hemoglobin oxidative degradation through H_2_O_2_ (Rifkind *et al*., 2003), have shown that nitrite interact with the Fe(II) heme, while the removal of oxygen is necessary for a direct reaction of nitrite with the heme-iron without oxygen bound. The distal histidine is required to protonate the nitrite as it binds (Ranghino *et al*., 2000), resulting in the cleavage of a hydroxide ion and the formation of the nitrosonium cation bound to the Fe(II) heme. The nitrosonium group is a stronger oxidizing agent than oxygen. As the oxygen pressure is reducing, nitric oxide (NO) production dominates and the NO is more stable, facilitating direct reactions involving NO but in the presence of oxygen, NO is unstable and exists in the form of N_2_O_3_ and nitrite (Rifkind *et al*., 2003). The production of labile bioactive NO in the red cell has been attributed to a redox reaction between nitrite produced in the plasma and hemoglobin (Rifkind *et al*., 2003).

The cascade begins with the peroxidase reaction which involve H_2_O_2_ with Fe(II) hemoglobin to produce the highly reactive Fe(IV) ferrylhemoglobin (ferrylHb), followed by the reaction of a second molecule of H_2_O_2_ with ferrylHb to produce a superoxide radical, which can attack the porphyrin before it escapes from the heme pocket (Nagababu and Rifkind, 2000). The initial damaged heme facilitates a change in the tetragonal geometry around the heme resulting in the formation of a high-spin rhombic heme complex (Nagababu *et al*., 2002). Gradual increase has been observed in the concentration of two fluorescent products, at 10 min for the 465 nm fluorescent product (excitation at 321 nm) and at above 20 min for the 525 nm fluorescent product (excitation of 460 nm) (Nagababu and Rifkind, 2000; 1998).

Another study has shown that deoxygenation enhanced the intensity of fluorescence by about 35 and 25% for the 465 nm emission band and the 525 nm emission band respectively, and the binding of carbon monoxide to Hb resulted in inhibition of about 95% of the fluorescence in both bands while peroxidase substrates such as ABTS and *o*-dianisidine completely suppressed the formation of both fluorescent products (Nagababu and Rifkind, 2000). Also, the distinctive roles of catalase and glutathione peroxidase occur at the initial steps of the degradation process and do not react with rhombic heme or fluorescent degradation products (Nagababu *et al*., 2003). It was reported that heme degradation was inhibited by the addition of catalase which removed hydrogen peroxide even after the maximal level of ferrylHb was reached (Nagababu and Rifkind, 2000). Additionally, study has shown that the formation of the heme fluorescent products during *in vivo* study involves oxidative stress (Eisinger *et al*., 1985). The fundamental features of chemiluminescent reaction include spontaneous less exothermic reaction that favors formation of reaction intermediate or product compound in the excited electronic state with emission of light, and its deactivation to the ground energy state (Kai *et al*., 2001).

## CONCLUSION

This study has provided a novel insight on the possible analogues of luminol present in the existing lead-like chemical database. The results have shown that blood artifacts can come other domestic organisms that are present in the human environment. The possible amino acid residues of hemoglobin that may be involved in the oxidative chemiluminescent reaction are also identified. Furthermore, improved mechanism of chemiluminescence of luminol in relation to heme oxidative degradation in the presence of hydrogen peroxide and alkaline medium was proposed. This will broader the possible applications of luminol with careful development of new methodologies.

## Competing interests

The author declares that there is no competing interest.

